# Direct and culture-independent detection of low-abundant *Clostridioides difficile* in environmental DNA via PCR

**DOI:** 10.1101/2023.07.24.550392

**Authors:** Miriam A. Schüler, Dominik Schneider, Anja Poehlein, Rolf Daniel

## Abstract

*Clostridioides difficile* represents a major burden to public health. As a well-known nosocomial pathogen whose occurrence is highly associated with antibiotic treatment, most examined *C. difficile* strains originated from clinical specimen and were isolated under selective conditions emplyoing antibiotics. This suggests a significant bias among analysed *C. difficile* strains, which impedes a holistic view on this pathogen. In order to support extensive isolation of *C. difficile* strains from environmental samples, we designed a detection PCR that targets the *hpdBCA* operon and thereby identifies low abundances of *C. difficile* in environmental samples. Amplicon-based analyses of diverse environmental samples demonstrated that the designed PCR is highly specific for *C. difficile* and successfully detected *C. difficile* despite its absence in general 16S rRNA gene-based detection strategies. Further analyses revealed the potential of the *hpdBCA* detection PCR sequence for initial phylogenetic classification, which allows assessing *C. difficile* diversity in environmental samples via amplicon sequencing. Our findings furthermore showed that *C. difficile* strains isolated under antibiotic treatment from environmental samples were originally dominated by other strains according to detection PCR amplicon results. This provided evidence for selective cultivation of under-represented but antibiotic-resistant isolates. Thereby, we revealed a substantial bias in *C. difficile* isolation and research.

**Importance:** *Clostridioides difficile* is mainly responsible for hospital-acquired infections after antibiotic treatment with serious morbidity and mortality worldwide. Research on this pathogen and its virulence focused on bacterial isolation from clinical specimen under antibiotic treatment, which implies a substantial bias in isolated strains. Comprehensive studies however require an unbiased strain collection, which is accomplished by isolation of *C. difficile* from diverse environmental samples and avoiding antibiotic-based enrichment strategies. Thus, isolation can significantly benefit from our *C. difficile*-specific detection PCR, which rapidly verifies *C. difficile* presence in environmental samples and further allows estimation of the *C. difficile* diversity by using NGS.

## Introduction

*Clostridioides difficile* (formerly *Clostridium difficile*) is a widely known pathogen, responsible for the majority of nosocomial infections worldwide with significant morbidity and mortality, especially after preceding antibiotic treatment (1). Symptom severity of a *C. difficile* infection not only depends on the individual physiological constitution but also on the infecting strain, as *C. difficile* virulence differs within the species (2, 3). Diverse characteristics of *C. difficile* are under research to elucidate its multifaceted pathogenicity (4). Thus, comprehensive investigations of the species *C. difficile* are essential to pinpoint the different characteristics of this pathogen. Most *C. difficile* strains isolated and examined to date originated from clinical specimens in the context of infections. However, this bacterium also resides in the intestinal microflora of healthy individuals and different animal species as well as environmental habitats like soil and water reservoirs (5). Consequently, there is a bias towards clinical-associated strains in *C. difficile* research (6, 7). Some researchers indeed already isolated *C. difficile* from the environment and thereby obtained multiple strains from one sample, including novel sequence types or even cryptic *C. difficile* strains that are classified as different genomospecies *Clostridioides* sp. by now (5, 6). However, further isolations from diverse habitats would contribute to a holistic view on this pathogen. Those broad-ranging isolation attempts would be promoted by initial assessment of environmental samples to harbor *C. difficile* before elaborate and time-consuming isolation experiments are performed. Common processes for *C. difficile* isolation last days up to few weeks and start with antibiotic-based cultivation to select for this presumably low-abundant species in complex environmental microbial communities (6). Selective cultivation is followed by sub-culturing and identification of isolates via e.g., MALDI-TOF, enzymatic immunoassays or 16S rRNA gene sequencing (5, 8, 9). The initial selective cultivation might eliminate environmental *C. difficile* strains that do not harbor the respective antibiotic resistance. For instance, a review from 2016 (10) summarized data from diverse studies on antibiotic resistances among clinical *C. difficile* isolates and showed that not all strains were resistant to the applied antibiotics, e.g. to cefoxitin that is used in *C. difficile* isolation attempts (11).

In this context, we designed a *C. difficile*-specific detection PCR for direct application on environmental DNA (metagenomic DNA, mgDNA), that produces a distinct amplicon upon *C. difficile* presence – even for an abundance below detection levels in general 16S rRNA gene amplicon-based community analysis (12). To our knowledge, this type of culture-independent detection directly performed on environmental DNA has not been established in *C. difficile* research so far. Previous PCR approaches for *C. difficile* detection worked on single isolates and pre-enriched samples (13–15) or targeted the toxin genes, resulting in strains lacking these genes or possessing different gene sequences are not detected (6, 16). PCR specificity was assessed by next-generation-sequencing (NGS) of amplicons that were produced by detection PCR on diverse mgDNA samples to prove *C. difficile* origin. Further, PCR sensitivity was demonstrated by comparison to the amplicon-based 16S rRNA gene analysis. Besides establishing and demonstrating successful performance of our PCR on diverse environmental samples, we examined the detection sequence as potential phylogenetic determinant via average-nucleotide-identity analysis. The most common methods for phylogenetic classifications of *C. difficile* strains are ribotyping and multilocus sequence typing (MLST) (17, 18). The former one involves amplification of the 16S-23S rRNA gene intergenic spacer region, which results in a genotype-specific pattern. This pattern has to be compared to already known ribotypes for identification, which is not feasible for all laboratories. Moreover, ribotypes can be polyphyletic and include various STs (19). The principle of MLST is based on sequence alleles of seven housekeeping genes, which can be determined via PCR and sequence analysis of all seven genes or whole-genome sequencing. Allele combinations are designated via the MLST database (http://pubmlst.org/) to sequence types (ST), which further group into monophyletic clades of C1 to C5 in the narrower sense of *C. difficile*, and additionally of cryptic clades C-I to C-V (6, 20). As such, MLST altogether exceeds ribotyping in phylogenetic classification of *C. difficile* strains. In contrast to MLST, phylogenetic classification with the here described detection PCR involves less effort and allows a superficial estimate of *C. difficile* strain diversity in environmental samples, which is so far not established in *C. difficile* research.

## Materials and methods

### *C. difficile* detection PCR target and primer design

To establish a *C. difficile*-specific detection PCR, literature research for species-specific genes or sequences were conducted and yielded amongst others the *hpdBCA* operon as potential target (21, 22). This operon encodes the *p*-hydroxyphenylacetate decarboxylase that catalyzes the production of *para*-cresol, an uncommon metabolic trait amongst bacteria (23).

The *hpdBCA*-nucleotide sequence was confirmed to be entirely present in all complete, unique *C. difficile* genomes deposited at the National Center for Biotechnology Information (NCBI (24); assessed 30 August 2022) using BLAST+ (v2.12.0) (25) with default parameters and the *hpdBCA*-sequence of *C. difficile* strain CD630 (NCBI accession CP010905.1, locus tags CDIF630_00272-00274) as query. A BLAST analysis (26) in megablast mode of the CD630 operon was done as well to check for general and significant occurrence of this sequence.

The primer design focused on several criteria: (I) no binding to possible non-*C. difficile* sequences while (II) binding to all *C. difficile* strains and additionally including so-called cryptic ones. These cryptic *Clostridioides* are closest related to *C. difficile* (6, 7) and further show >99.87% 16S rRNA gene sequence similarity to *C. difficile*, which would even classify them as same species (27). (III) Yielding a PCR product of <500 bp length for amplicon analysis with NGS. (IV) Fulfilling standard criteria such as similar melting temperature, GC clamp, no secondary structure and no self-/cross-dimer formation (http://www.premierbiosoft.com/tech_notes/PCR_Primer_Design.html).

(I) The *hpdBCA*-sequence of CD630 (3,913 bp) was used as reference in a blastn analysis excluding *C. difficile* (taxid 1496) to retrieve all similar non-*C. difficile* sequences. The alignment of these sequences to the CD630 reference was inspected using the NCBI Multiple Sequence Alignment Viewer (MSA Viewer, v1.22.0, https://www.ncbi.nlm.nih.gov/tools/msaviewer/) for promising primer regions that show least similarity between the CD630 and the non-*C. difficile* sequences, with special focus on the 3’-primer end as being the crucial part for amplification (28) (Fig. S1). (II) Potential primer regions from step (I) were examined regarding conservation across *C. difficile*. Therefore, all aligned *hpdBCA* sequences from the previous megablast analysis against complete *C. difficile* genomes at chromosome level were downloaded and aligned in AliView v1.26 (29). (III) Corresponding forward/reverse primers for defined primer candidates were searched to amplify a 400-500 bp sequence while fulfilling criteria (I) and (II). Standard primer criteria such as GC clamp, similar melting temperature and no self- or primer-dimer formation (IV) were examined with the online tools of ThermoFisher Scientific (Multiple Primer Analyzer, https://www.thermofisher.com/de/de/home/brands/thermo-scientific/molecular-biology/molecular-biology-learning-center/molecular-biology-resource-library/thermo-scientific-web-tools/multiple-primer-analyzer.html) and Sigma-Aldrich (OligoEvaluator http://www.oligoevaluator.com/LoginServlet). Performance comparison of all potential primers resulted in the final pair of CDIF_cresol_3F (forward 5’-GAAAAGGTGGGTTCCATATTCAATATAATG) and CDIF_cresol_3R (reverse 5’-CCTTCTAATTGCTTTTGACTACTCATTAAACAC). Both primers aligned to all analyzed *C. difficile* strains with 100% coverage and predominantly 100% percent identity except for C5 strains with one mismatching nucleotide (forward 5’-position: 8, reverse 5’-position: 11; see also Fig. S2), but still allowing efficient primer annealing. The cryptic *Clostridioides* strains showed also a coverage of 100% of the primers but percent identities lay between ∼91% and ∼96%, corresponding to one to four mismatching nucleotides.

### Bioinformatic analyses of detection PCR primers and sequence

#### *C. difficile* specificity analysis

The detection PCR primers were examined *in silico* for *C. difficile*-specificity by blastn analysis with default settings excluding *C. difficile* (taxid 1496) (done on 15 September 2022). *C. difficile*-specificity of the corresponding detection PCR product was likewise assessed by blastn analysis (done on 15 September 2022), using the sequence retrieved from CD630 via *in silico* PCR. The primer sequences were thereby omitted so that only the 401 bp-long sequence between the primers was used as BLAST query.

#### Detection PCR sequence as phylogenetic marker

The 401 bp-long detection sequence was further analyzed for its potential to phylogenetically classify *C. difficile* isolates. Phylogenetic examinations at whole genome and detection sequence level were performed using all complete *C. difficile* genomes available at NCBI by average-nucleotide-identity (ANI) calculations with PYANI v0.2.11 (30)(31) in default settings and MUMmer3 (32) for sequence alignment (ANIm). ANIm results were visualized in Rstudio with the gplots-implemented tool heatmap.2. Phylogenetic assignment of the analyzed strains in form of MLST determination was performed with FastMLST v0.0.15 (33).

Alignment of unique detection sequences including the primers amongst the above analyzed complete *C. difficile* genomes corresponding to specific MLST types/clades were visualized in Jalview v. 2.11.2.0 (34) for comparison of MLST/clade-associated nucleotide deviations.

### Isolation of mgDNA from environmental samples

Diverse environmental samples (n=46, Table 1, detailed information in Table S1) such as soil, compost or faeces, of which the majority had been in contact to animals, were collected in sterile canonical falcon tubes and stored at 4 °C upon arrival in the laboratory. The mgDNA was extracted with the DNeasy PowerSoil Pro kit (Qiagen) according to protocol. DNA concentration was determined with the Qubit 3.0 Fluorometer (Life Technologies) using the BR dsDNA assay kit and DNA purity was assessed via NanoDrop ND-1000 (Peqlab Biotechnologie GmbH) measurement.

**Table 1.** Environmental samples used in this study with sample ID and description.

| ID | Description | ID | Description |
| --- | --- | --- | --- |
| A1 | Horse turd | P2 | Horse turd (mother to J2) |
| B1 | Sludge biogas fermenter | P3 | Horse turd |
| D1 | Muddy trickle | PA | Horse turd |
| Goe1 | Compost – disposal company | TS10a | Wild boars wallow |
| Goe2 | Mud dump | TS10b | Wild boars dung pile |
| J1 | Meconium foal 1 | TS10c | Wild boars dung |
| J2 | Meconium foal 2 | TS1a | Donkeys dung pile |
| J3 | Horse turd foal 3 | TS1b | Donkeys dung |
| J4 | Horse turd foal 4 | TS2ab | Red cattle dung pile + dung |
| J5 | Horse turd foal 5 | TS3 | Mud from outdoor area farm |
| J6 | Horse turd foal 6 | TS4ab | German saddlebacks dung pile + dung |
| J7 | Horse turd foal 7 | TS5a | Exmoor ponies dung pile |
| K1 | Wild boars dung | TS5b | Exmoor ponies dung |
| K2 | Wild boars dung pile | TS6 | Mud from swampy region (outside enclosure) |
| K3 | Wild boars wallow | TS6a | Mud in fallow deer area |
| K4 | Fallow deer dung | TS7a | Reindeer dung pile |
| K5 | Fallow deer dung pile | TS7b | Reindeer dung |
| MA | Horse dung | TS8a | Woolly pigs wallow |
| MG1 | Slurry | TS8bc | Woolly pigs dung pile + dung |
| MG2 | Dung pile calf stable | TS9a | Pot-bellied pigs wallow |
| MG3 | Mud from cow run | TS9b | Pot-bellied pigs dung pile |
| MG4 | Sludge from biogas fermenter | TS9c | Pot-bellied pigs dung |
| P1 | Horse turd | TSA | Muddy trickle |

#### Detection PCR on genomic DNA (g-PCR)

Optimal detection PCR conditions were initially assessed on genomic DNA of in-house *C. difficile* strains belonging to different MLSTs/clades (ST1/C2, ST3/C1, ST8/C1, ST11/C5, ST340/C-III). PCR was performed with DreamTaq polymerase (Thermofisher Scientific) according to manufacturer’s recommendations using 0.2 mM of each primer and 50 ng DNA template per 50 µl PCR reaction mixture. Established PCR cycling conditions comprised initial denaturation at 95 °C for 3 min, followed by 28 cycles of denaturation at 95 °C for 30 s, primer annealing at 61 °C for 30 s and extension at 72 °C for 40 s, and final extension at 72 °C for 5 min.

#### Detection PCR on metagenomic DNA (mg-PCR)

The previously established g-PCR parameters were adapted for detection on mgDNA. Considering that mgDNA is a diverse mixture of genetic material with presumably minor abundance of *C. difficile* DNA, we aimed to support primer annealing at a low abundant template. Adaptations included doubled DNA template amount of 100 ng per 50 µl PCR reaction mixture, increased primer concentration of 0.3 mM, and modified cycling conditions with 35 cycles comprising a specific time decrement program during primer annealing for 90 s that decreases by 2 s each cycle.

#### Analysis of bacterial community composition via 16S rRNA gene amplicon sequencing

For sensitivity assessment of the detection PCR compared to the common investigation of bacterial community composition, we conducted a 16S rRNA gene analysis of all 46 samples (12). Amplification of the V3 to V4 region of the 16S rRNA gene was performed in triplicates with Phusion™ High-Fidelity DNA polymerase (Thermofisher Scientific) as stated in the protocol using GC buffer, 5% DMSO, 0.2 mM of each primer S-D-Bact-0341-b-S-17/S-D-Bact-0785-a-A-21(12) with attached Illumina amplicon adapter overhang sequences (35) and 25 ng mgDNA template per 50 µl PCR reaction mixture. PCR triplicates were pooled equimolar, purified, and sequenced as described by Berkelmann et al. (36).

Illumina raw-reads were processed with an in-house pipeline as follows and thereby parallelized with GNU parallel 20190322 (37): raw-read quality filtering with fastp v0.23.2 (38) (extended default settings as in (39)), merging of paired-end reads with PEAR v0.9.11 (40), and clipping of primer sequences using cutadapt v3.2 (41). VSEARCH v2.12.0 (42) was used to perform size sorting and filtering (>300 bp), read dereplication, denoising with UNOISE3 (minimum abundance option: minsize 8) (43), chimera removal with UCHIME3 (44) *de novo* and reference-based against the SILVA SSU 138.1 NR database (45), and final read mapping to amplicon sequence variants (ASV) at 100% identity to create an abundance table. ASV taxonomies were assigend with BLAST 2.9.0+ (46) using the SILVA SSU 138.1 NR database with a 70% identity cut-off, and added to the abundance table with BIOM tools v1.0 (47). Final amplicon data was further processed in R by applying taxonomic boundaries according to sequence identities as defined by Yarza et al. (48), and removing spurious sequences by applying a 0.25% cutoff filter on each sample according to Reitmeier et al. (49) before visualization with the R package ggplot2 (50).

Alpha diversity was calculated and visualized with the R packages ampvis2 (51) and ggplot2 (50) using “amp_alphadiv” and “amp_ordinate” functions.

### Sequencing of detection PCR amplicons

Sequencing of PCR products from positive detections on diverse mgDNA samples was performed to verify *C. difficile* origin and, thus, assess PCR specificity and sensitivity. Sequencing was done with *C. difficile* isolates obtained from the analyzed environmental samples as indication and control for correct processing of amplicon data. For Illumina sequencing, the Illumina amplicon adapter overhang sequences (35) were attached to the 5’-end of the detection primers. Amplicon PCR was performed in triplicates with Phusion™ High-Fidelity DNA polymerase (Thermofisher Scientific) according to manufacturer’s indications using GC buffer and no DMSO. The PCR parameters were used as determined in the above-mentioned mg-PCR protocol, including thermocycling conditions. All corresponding replicates were pooled equimolar before purification with the GeneRead Size Selection kit (Qiagen) as recommended by the manufacturer, including two successive purification rounds with repeated elution for highest amplicon recovery and elimination of primer-dimers. First and second purification round were eluted in 90 µl and 15 µl EB buffer, respectively. DNA concentration was determined in the Qubit 3.0 Fluorometer (Life Technologies) using the HS dsDNA assay kit. Amplicon quality was assessed employing a Bioanalyzer and the DNA 1000 kit as recommended by the manufacturer (Agilent).

Raw reads were processed as described for 16S rRNA gene amplicon sequencing with few adaptations. The ASV size filter was set to ≥380 bp and chimeras were removed *de novo*. ASV taxonomies were assigned with BLAST against the NCBI BLAST database (as from 5 October 2022). MLST assignment of each ASV based on its BLAST identity (Genbank accession) was performed using the PubMLST database (52).

Final amplicon data was further processed in R. Based on data situation and according to Reitmeier et al. (49), an appropriate cutoff filter of 0.25% as employed in 16S-amplicon data processing was finally applied, and data was visualized with the R package ggplot2 (50).

### Data records

The data was deposited at the National Center for Biotechnology Information and can be found under the BioProject accession numbers PRJNA991070, PRJNA991304, and PRJNA991297. These BioProjects contain the raw sequences of 16S rRNA gene amplicons (SRR25194157-SRR25194209) and of detection PCR amplicons (SRR25186671-SRR25186686 and SRR25132251-SRR25132254).

## Results

### Initial specificity assessment of *C. difficile* detection PCR

We initially assessed the *C. difficile* specificity of the detection PCR with bioinformatics examinations using BLAST. First, all unique and complete *C. difficile* genomes at chromosome level – 134 in total – were verified to contain the entire *hpdBCA*-sequence by BLAST+ alignment against the CD630 operon, which showed overall 100% query coverage and above 97% percent identity. A further BLAST megablast analysis of the CD630 operon was performed to check for significant non-*C. difficile* matches. Only significant matches against *C. difficile* entries and five genomes belonging to the cryptic *Clostridioides* strains were detected. The BLAST alignments against these *Clostridioides* sp. strains showed above 99% query coverage with over 92% percent identity to the CD630 operon.

All unique *C. difficile* genomes and the five *Clostridioides* sp. strains – 139 in total (Table S2) – were used for ANIm calculations.

The detection primers amplified a 464 bp-long part of the 3.9 kb *hpdBCA*-operon (position 2,525 to 2,988) that spanned across the entire *hpdC* gene and parts of genes *hpdB* and *hpdA*. The reverse primer covered *hpdC* and *hpdA*, which corresponded to an alignment gap in similar sequences in most non-*C. difficile* species, as determined by blastn *hpdBCA*-sequence alignment during primer design (Fig. S2). Consequently, the reverse primer was primarily responsible for *C. difficile*-specificity. However, two strains – *Clostridium diolis* (CP043998.1) and *Pelosinus sp.* strain UFO1 (CP008852.1) – did not show this alignment gap but possessed sufficient sequence deviation to rule out effective binding of the reverse primer in theory (Fig. S2). Detection mg-PCR on genomic DNA of these strains proved to be true negative and, thus, supported PCR specificity. Further, mg/g-PCR was verified to be positive on a self-isolated cryptic C-III strain so that successful detection of at least C-III cryptic *Clostridioides* was verified.

### Potential of the detection sequence *hpdBCA* as phylogenetic determinant

Heterogeneity of the detection PCR sequence and, thus, its potential for phylogenetic classification of *C. difficile* isolates, was examined by ANIm analysis (hereafter referred to as detANIm) in relation to a whole-genome ANIm (wgANIm) comprising all 139 genomes defined above (data of ANIm outputs in Table S3 and S4).

The wgANIm of the 139 NCBI strains (Fig. 1) exhibited clustering as already described for *C. difficile*, with a cluster-group comprising five distinct clusters and an overall ANI above 96% (species boundary) (7). Based on determined MLSTs (Table S2), each of the five clusters was assigend to clades C1 to C5. Intra- and inter-clade comparisons of mean, median, minimal and maximal ANIm values (Fig. 2) revealed that isolates within one clade shared ANI values above 99.24%. One isolate (ST963) could not be assigned to a clade with FastMLST, and did also not apparently cluster to a specific clade within the wgANIm. This was further visible by its inter-clade ANI values below 99% (Fig. 2), with highest similarity to C4. Subsequently, this isolate was treated as unclassified outlier. C1 was the biggest cluster, followed by C2, C5, C4 and finally C3. Additionally, C1 also showed highest heterogeneity among the five clades, indicated by a broader ANI range and apparent sub-clustering. C5 was least similar to C1 to C4 than these clades amongst themselves.

**Figure 1.**
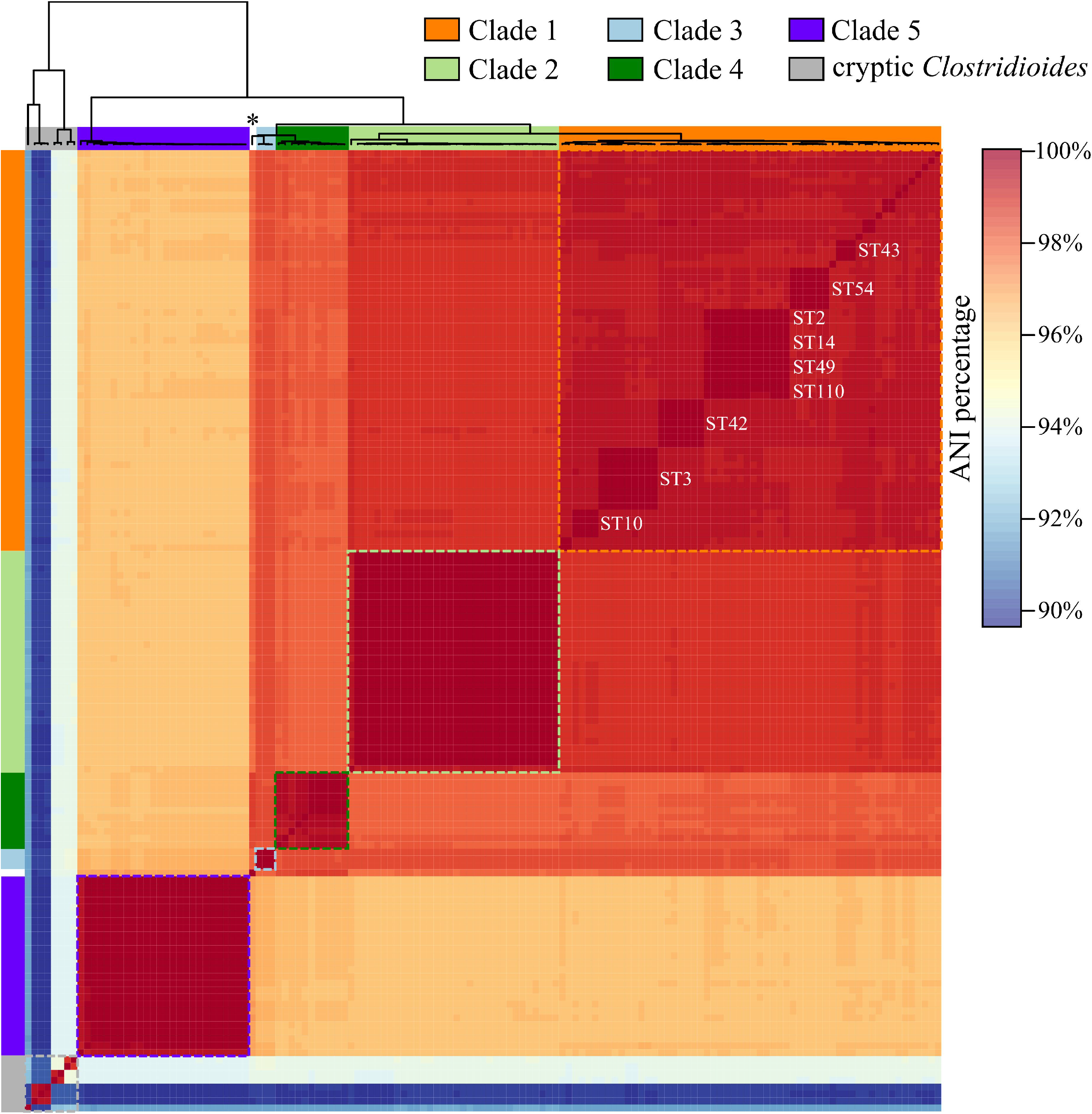
Heatmap of whole-genome ANIm analysis (wgANI). ANIm percentage values were visualized in heatmap.2 with corresponding dendrogram. Phylogenetic clades based on MLST determination are color-highlighted and MLST types are noted for C1 sub-clusters comprising more than three strains. The outlier strain ST963 is marked with * in the dendrogram.

**Figure 2.**
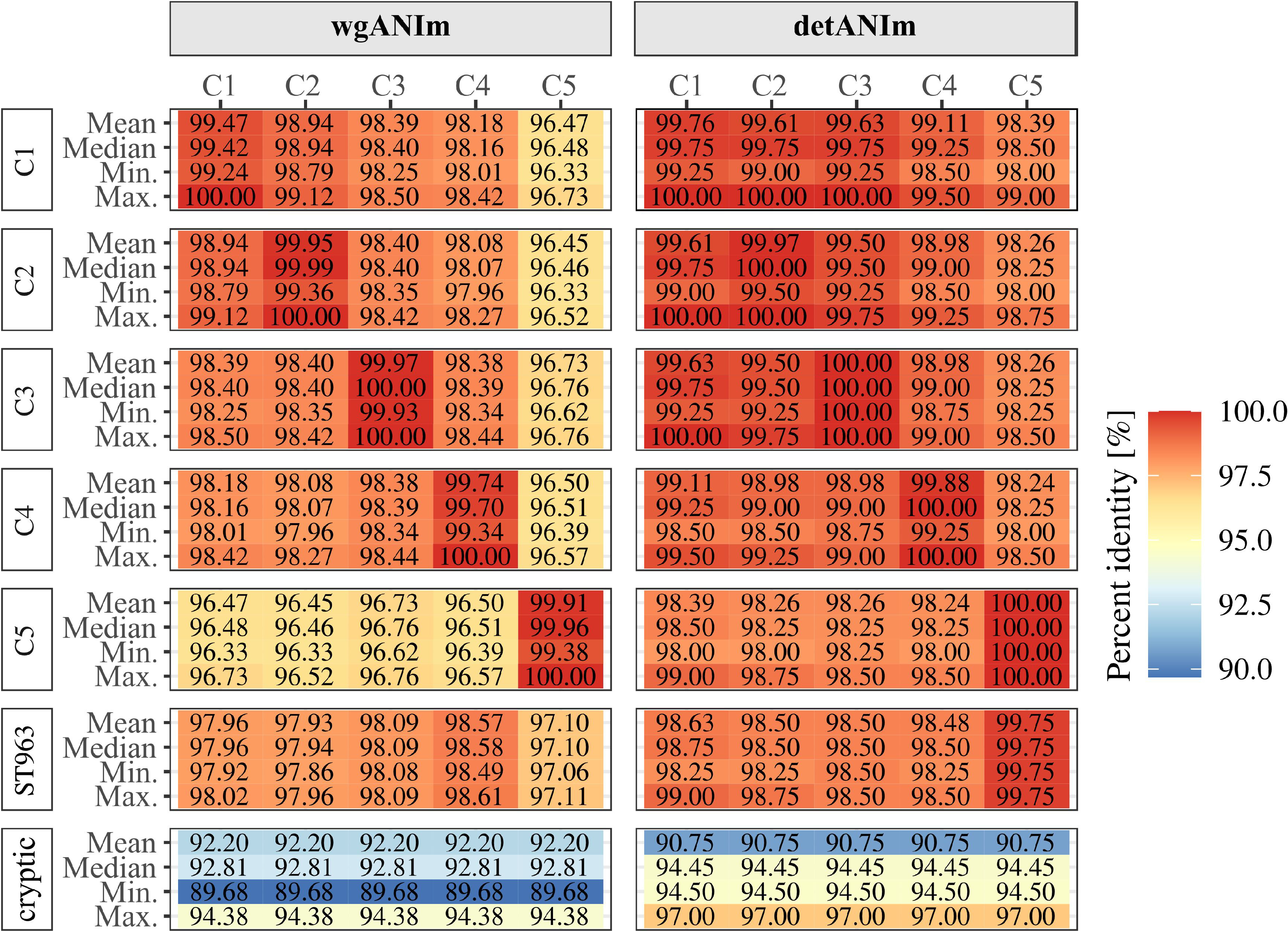
Clade-wise comparison of percentage identities for wgANIm and detANIm. Given are mean, median, minimal and maximal ANI percentages between two clades of C1-C5, outlier ST963 to each clade of C1-C5 or between all cryptic *Clostridioides sp.* to *C. difficile* (all five clades combined). Clade affiliation of an isolate was defined by MLST assignment using FastMLST (33).

In addition to clades C1 to C5, four further separate clusters were visible in the wgANIm, which represented the clades of cryptic *Clostridioides*. These clusters shared below 94.38% ANI to the five clades (Fig. 2) and comprised the five *Clostridioides* sp. (cryptic clades C-III and C-IV (6)) as well as three novel *C. difficile* strains (Cdiff_RT151_6, Cdiff_RT151_7, Cdiff_RT151_8), indicating that these strains belong to the cryptic *Clostridioides*. This was confirmed by further ANIm calculations with these strains against reference strains of cryptic clades C-I to C-V (6), which identified them as C-I and C-II strains, respectively.

The detANIm analysis revealed apparent clustering of specific sequence variants (Fig. 3), which resembled the wgANIm clustering. Alike the wgANIm, a large group of several clusters contained all *C. difficile* strains, while smaller separate clusters comprised the five *Clostridioides* sp. and the three presumable cryptic *Clostridioides* strains. These separate clusters shared below 97% ANI to *C. difficile* strains, while *C. difficile* detection sequences amongst C1 to C5 strains were at least 98% similar and often 100% identical within clusters (Fig. 2).

**Figure 3.**
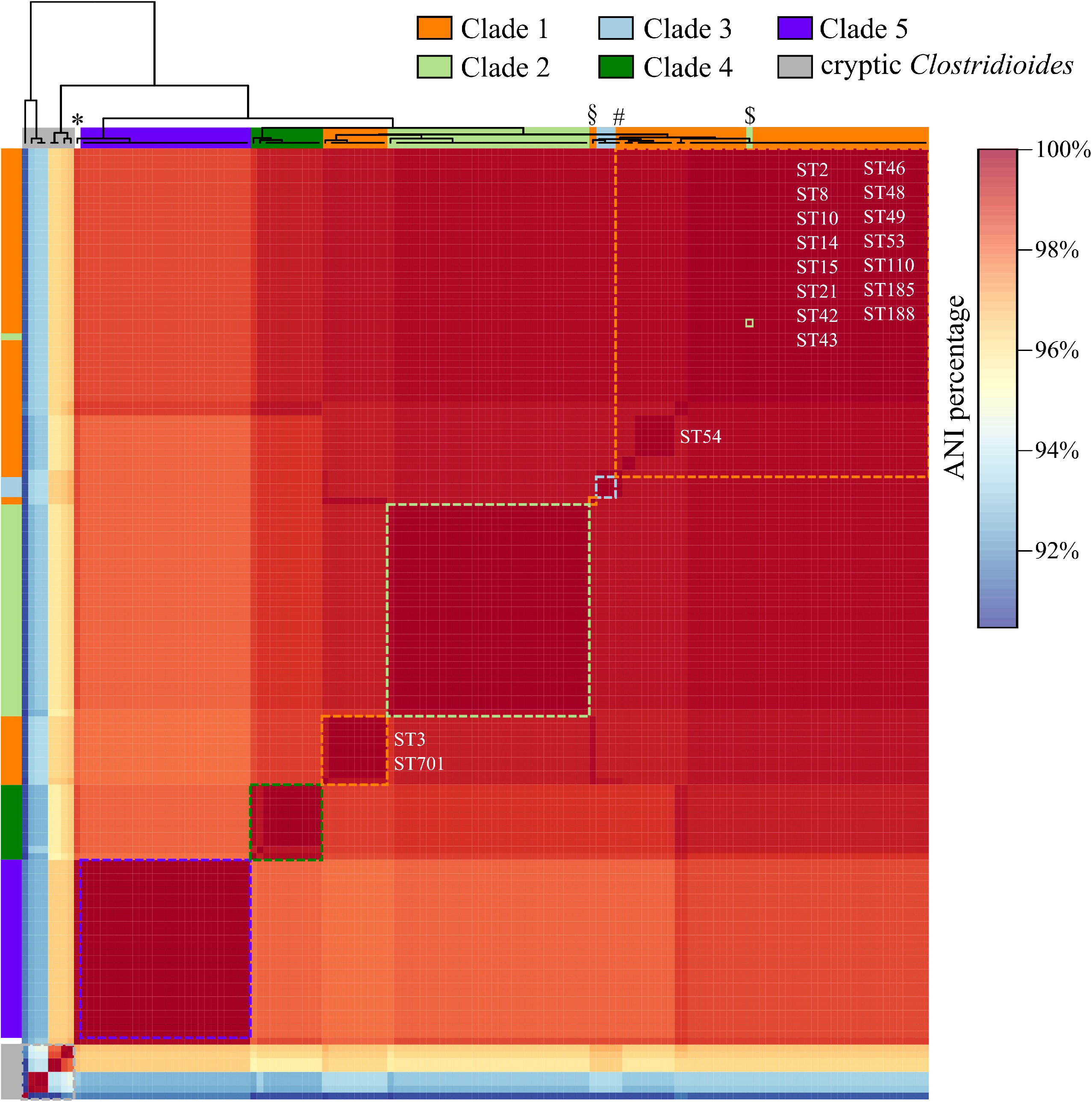
Heatmap of detection sequence ANIm analysis (detANIm). ANIm percentage values were visualized in heatmap.2 with corresponding dendrogram. Phylogenetic clades based on MLST determination are color-highlighted and MLST types are noted for C1 sub-clusters comprising more than three strains. Outliers are marked as follows in the dendrogram: strain ST963 with *, C1 outliers DSM 29632 and Z31 with § and #, C2 outlier Cd1 with $.

For detailed phylogenetic analysis of the detection sequence, previous MLST assignments of the genomes included in ANIm calculations were transferred to the corresponding detection sequences in the detANIm. This revealed a slightly different clustering or cluster positioning of the five clades. C1 sequences showed again highest heterogeneity (Fig. 2 and 3) but split into sub-clusters that partially shared 100% ANI and comprised various MLSTs. Further, sequences of two C1 strains (DSM 29632 and Z31) did not cluster in their correct clade or were even more similar to C3 sequences. Likewise, the wgANIm, detection sequences of C2 strains were closest related to those of C1 strains, with one outlying C2 strain (Cd1) even clustering within C1 sequences. This strain already attracted attention in the wgANIm by sharing ANI values to the other C2 strains below 99.40%, whereas all other C2 strains showed higher sequence similarity of at least 99.94%. All detection sequences of C3, C4 and C5 isolates clustered according their phylogenetic clade. The detection sequence of wgANIm-outlier strain ST963 with undefined MLST/clade and highest wgANIm similarity to C4 now clustered with C5. However, while all sequences within C5 were 100% identical, strain ST963 was distinguishable and only shared 99.75% ANI to all C5 isolates (Fig. 2).

Alignment of all 139 detection sequences revealed the unique nucleotide differences that represent distinct ANIm-clusters and MLSTs/clades (Fig. S3). All detection sequences used for ANIm analyses are listed in the supplementary Data File S1, with GenBank accession numbers of the corresponding *C. difficile* genomes as well as MLST/clade information as determined with FastMLST (33) or otherwise deduced from ANIm results. This file can be used for future ANI calculations or employed as BLAST+ database to examine one’s own detection sequences of interest.

Further apparent in Fig. S3, the primer regions are conserved amongst diverse *C. difficile* strains except for the already mentioned mismatch in the regions of C5 strains.

### Amplicon-based examination of detection PCR sensitivity and specificity

Detection PCR sensitivity was evaluated in comparison to the common 16S rRNA gene analysis. None of the 46 environmental samples contained the *C. difficile-*16S rRNA gene ASV in the final amplicon data after removing spurious sequences with the 0.25% cutoff filter (49), which eliminated the few present *C. difficile*-ASV reads (maximal 0.029% per sample) (Data File S2). Consequently, this analysis suggested the lack of *C. difficile*. However, this data based on the standardized 16S rRNA gene PCR with 25 cycles whereas the detection PCR involves 35. Thus, PCR sensitivities were not directly comparable. To rule out that *C. difficile* absence in 16S rRNA gene-amplicon data resulted from insufficient PCR cycling, 16S rRNA gene PCR was repeated with 35 cycles using specific environmental samples that possessed the *C. difficile*-ASV in the unfiltered data or from which we had isolated *C. difficile*. In this final amplicon data, only one out of eight analyzed samples contained reads corresponding to the *C. difficile*-ASV at all, and the abundance of 0.0134% lay again below the 0.25% filter cutoff (Data File S3). These findings supported that general 16S rRNA gene sequencing-based analyses suggest *C. difficile* absence in the environmental samples.

Further analyses of the 16S rRNA gene-amplicon data focused on community composition and diversity (Fig. S4-6). Bacterial abundance at genus was level diverse within and between the environmental samples (Fig. S4 and S5a) and similar composition at genus level was rather linked to sample type (Fig. S4 A) but not to detection PCR outcome (Fig. S4 B). Alpha diversity was assessed as ASV richness (Fig. S6). Most environmental samples showed a richness between 2,000 and 6,000 ASVs. Thereby, no correlation between ASV richness of a sample and the corresponding detection PCR result was apparent.

Detection PCR specificity was already supported experimentally by true negative results using genomic DNA of bacteria that were suspected to be false positive (*C. diolis* and *Pelosinus sp.* strain UFO1). For further confirming PCR specificity, NGS of detection PCR products was performed to examine the origin of amplicon sequences. First, the original detection on mgDNA was positive for 16/46 environmental samples with varying PCR product yield as seen by band strength on agarose gel, which might be an indicator for *C. difficile* abundance. The detection PCR for NGS purpose was performed using the 16 samples yielding positive results. Multiple reactions per PCR replicate were necessary for samples where low PCR product yield was observed in the original detection PCR. For assessing adequate processing of detection amplicon data, NGS was performed using control amplicons from four different *C. difficile* isolates that were isolated from three of the environmental samples, which should only give rise to one sequence. Indeed, final amplicon data for these isolates contained only one ASV each with 100% relative abundance. These ASVs matched the expected detection sequence for each isolate as determined by *in silico* PCR. The final amplicon data of all environmental samples contained only ASVs with *C. difficile* identity, proving that all detection PCR products originated from *C. difficile* DNA present in the mgDNA samples (Fig. 4).

**Figure 4.**
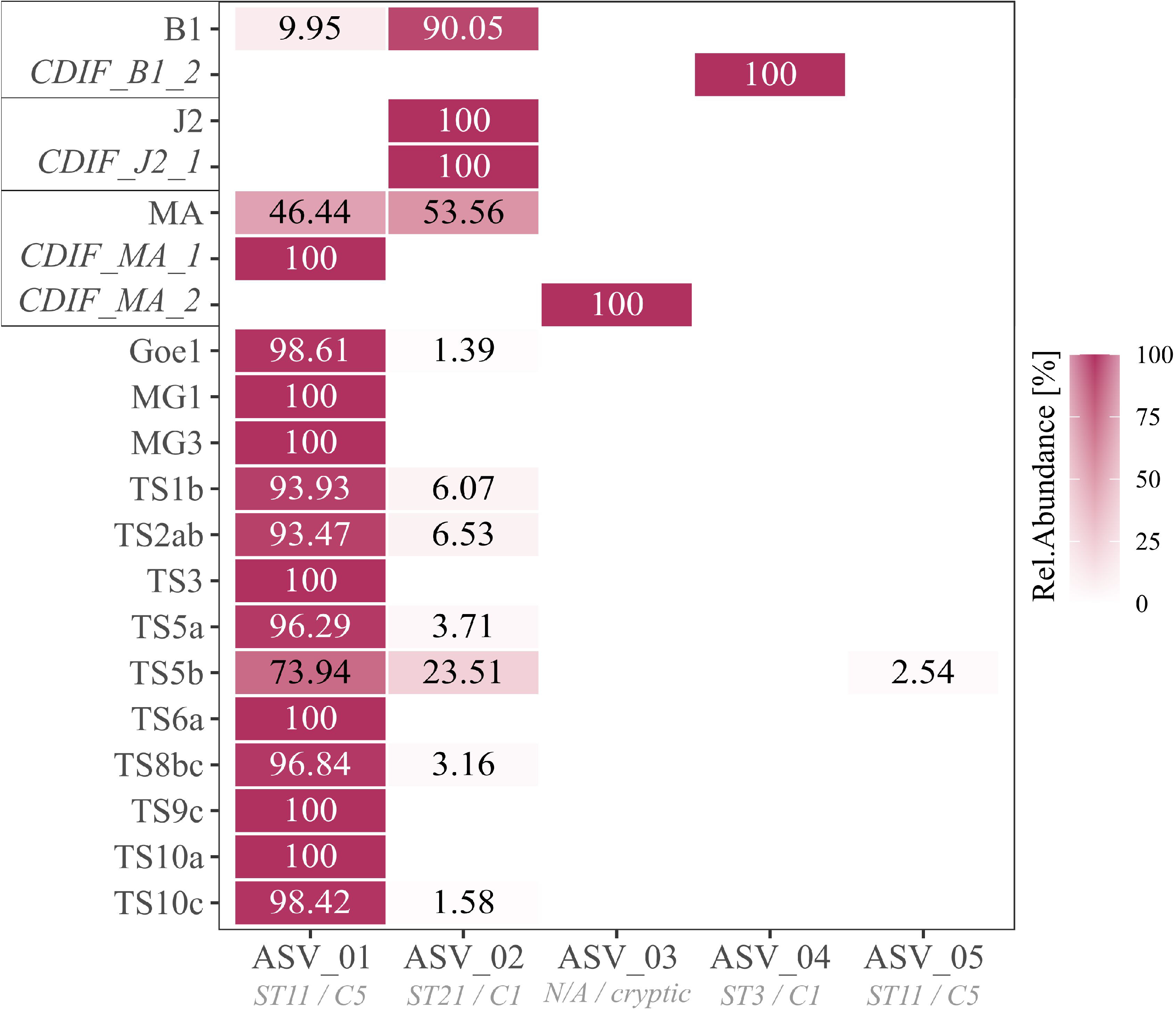
Heatmap of detection-ASV distribution and their relative abundances for selected samples. C. difficile isolates are in italic and depicted next to their original environmental sample. MLST assignment (ST / clade) of each ASV as described in Table 2 is additionally specified below each ASV. Percentage values of the relative abundances are noted for values >0%.

**Table 2.** BLAST results from amplicon data processing of the five ASVs shown in Fig. 4. Listed are BLAST hits with species and strain (Genbank accession), percentage identity, query coverage and E-value. Additionally, MLST/clade assignment for each strain is given as determined via PubMLST.org.

| ASV | BLAST hit | Identity (%) | Query coverage (%) | E-value | MLST | Clade |
| --- | --- | --- | --- | --- | --- | --- |
| 01 | *C. difficile* strain TW11-RT078<br>CP035499.1 | 100 | 100 | 0.0 | ST11 | C5 |
| 02 | *C. difficile* strain 020709<br>CP028529.1 | 100 | 100 | 0.0 | ST21 | C1 |
| 03 | *C. difficile* strain ST632<br>CP103804.1 | 98.005 | 100 | 0.0 | ST632 | cryptic<sup>a</sup> |
| 04 | *C. difficile* strain DSM 29745<br>CP019857.1 | 100 | 100 | 0.0 | ST3 | C1 |
| 05 | *C. difficile* strain TW11-RT078<br>CP035499.1 | 98.504 | 100 | 0.0 | ST11 | C5 |

<sup>a</sup>no specific cryptic clade could be assigned to CP103804.1, highest similarity was found to C-III with a percentage identity of 95% - 96% based on ANIm analysis against reference strains for cryptic clades C-I to C-V selected from (6).

The environmental samples comprised five different ASVs representing four sequence types within three phylogenetic clades, amongst others a cryptic clade (Fig. 4, Table 2). From the 16 mgDNA samples, seven samples contained one, eight samples two, and sample TS5b comprised even three *C. difficile*-affiliated ASVs. The most prevalent ASV_01, classified as ST11/C5 sequence, was present in all mgDNA samples except of sample J2. The second most abundant ASV_02 with determined ST21/C1 identity was present in ten mgDNA samples with an abundance ranging from 1.39% to 90.05%. These two ASVs were present in control PCRs with DNA from *C. difficile* isolates MA_1 (ST11/C5) and J2_1 (ST8/C1), coinciding with ASV presence in their original environmental sample MA and J2. Contrary, the cryptic isolate MA_2 (ST340/C-III) correctly exhibited ASV_03, which was however not observed in the corresponding environmental sample MA used for isolation. Environmental sample B1 also comprised the two most abundant ASV_01 and ASV_02. Remarkably, *C. difficile* strain B1_2 that was isolated from sample B1 did not possess any of these ASVs but the ST3/C1 sequence ASV_04. It should be noted here that both isolates MA_2 and B1_2, which were not observed in the detection amplicons of their respective environmental sample, were isolated with antibiotic-based cultivation according to Dharmasena & Jiang (8).

The other environmental samples (Goe1 – TS10c in Fig. 4) were dominated by ASV_01 with at least 93.47% relative abundance, often accompanied by the second ASV_02 with maximal 6.53% abundance. The only exception was the afore-mentioned sample TS5b, in which ASV_01 comprised 73.94%, ASV_02 23.51% and the third ASV_05 2.54%. ASV_05 was as ASV_02 identified as ST11/C5 sequence. However, although matching best to C5 sequences, the similarity of ASV_05 to the C5-BLAST reference of 98.504% lay below our previously determined intra-clade detANI value (100%) of C5 strains (Fig. 2).

## Discussion

We established a PCR that specifically detects the global pathogen *C. difficile* in environmental DNA, even in low abundance. Thus, environmental samples can be evaluated for *C. difficile* presence, which in turn can support *C. difficile* isolation from various environments and thereby contributes to holistic investigations of this significant pathogen. In addition to detection purposes, the detection PCR allows to initially classify the phylogenetic affiliation of *C. difficile* isolates or to assess *C. difficile* diversity present environmental samples.

The sensitivity of our detection PCR was validated in comparison to the commonly used amplicon-based 16S rRNA gene analysis. This method did not show *C. difficile* in various environmental DNA samples, meaning the abundance of *C. difficile* 16S rRNA gene amplicons was below the filter cutoff during amplicon processing or not present at all. Even increased PCR cycling in 16S rRNA gene-amplicon production as adjustment to the higher cycle amount within the detection PCR did not reveal *C. difficile* presence. Contrary, our detection PCR was positive for one third of the environmental samples. As such, the PCR was proven to outperform the 16S rRNA gene approach in detection of low-abundant *C. difficile,* even without prior enrichment as done in other studies (15).

Detection PCR specificity was evaluated by NGS of detection amplicons from diverse environmental samples. The bacterial community of these samples was divergent as shown by 16S rRNA gene analysis. Sequencing of detection amplicons obtained from those diverse environmental samples verified high *C. difficile-*specificity of the detection PCR, as only *C. difficile*-ASVs were found. With this proof of specificity, the detection PCR can be used beyond the detection purpose in environmental DNA and quickly verify newly isolated strains as *C. difficile* in a simple colony PCR. This identification procedure is a fast and easy alternative to common methods such as MALDI-TOF, 16S rRNA gene sequencing or enzymatic assays (5, 8, 9). We also examined the detection sequence itself as potential phylogenetic determinant via ANIm analyses with regard to a whole-genome ANIm. ANIm clusters in combination with MLST assignment showed that clade sizes and MLST type distribution among our genome dataset matched the holistic, taxonomic investigation by Knight et al. (7), supporting that the dataset adequately represented the species *C. difficile.* ANIm clusters observed at whole-genome level were mostly identified in the detANIm, which showed that the detection sequence reflects phylogenetic information to some extent. While the detection sequence allowed correct clade assignment for almost all 139 analyzed strains except of the four described outliers, it could not distinguish between closely related STs. Thus, a complete phylogenetic classification with distinct MLST assignment requires detailed MLST analysis. Nevertheless, the detection PCR not only allows identification of *C. difficile* isolates with a positive PCR result but also enables an initial phylogenetic classification of the isolate by sequencing the recovered PCR product.

Further, we investigated the NGS detection amplicons from environmental DNA samples and predominantly found sequences that match *C. difficile* ST11/C5 and ST21/C1 strains. Prevalence of C5 strains is linked to various animal species such as horses, pigs and cattle (53). These observations coincide with our results of prevalent C5 strains in the analyzed environmental samples, of which the majority were in previous contact with the afore-mentioned animals. Prevalent sequences with C1-association in the environmental samples in turn reflect the global occurrence of C1 strains in both animals and humans, and are in this context also of interest concerning transmission of potentially toxinogenic strains (20, 54, 55). Noteworthy, in the light of our detANIm analysis, the sequence of ST21/C1 is not unique for this sequence type but shares 100% similarity to sequences of further C1 sequence types. This was demonstrated by the environmental sample J2 and the corresponding isolate J2_1 for which the detected ASV_02 was assigned to ST21/C1, albeit strain J2_1 belonging to ST8/C1. Consequently, detection ASVs with determined ST21/C1 identity might belong to another C1-ST sequence as well and even the presence of multiple C1 strains in the environmental sample can be considered.

We also performed the detection amplicon sequencing on *C. difficile* strains that originated from the environmental samples, and checked for ASV accordance between the isolates and their corresponding environmental sample. The two analyzed strains MA_2 and B1_2 and their ASVs did not accord with ASV presence in the corresponding environmental data, which instead contained ASVs of two different *C. difficile* strains. This implied that the isolated strains were insufficient abundant in the environmental sample to be identified by the detection PCR, contrary to those strains that showed up in the detection amplicons. In that regard, the successful isolation of these strains despite their underrepresentation indicated a selective cultivation of these underrepresented strains, in contrast to the other more abundant strains. Such selective cultivation can be performed for instance with antibiotic treatment. Indeed, we obtained these *C. difficile* isolates with antibiotic-based cultivation using moxalactam norfloxacin, an established supplement for selective *C. difficile* isolation (8, 56). In conclusion, these results suggest an abundance shift of *C. difficile* strains in environmental samples by applying selective antibiotic cultivation procedures with environmental samples as starting material, which supports our hypothesis to miss some strains during antibiotic-based isolation approaches. Thus, for assessing *C. difficile* in environmental samples, a combination of both strategies is advisable.

## Acknowledgements

We thank the laboratory of Prof. Dr. Adams at the Department of Biochemistry and Molecular Biology, University of Georgia, Athens, GA 30602, USA for providing the genomic DNA of *Pelosinus sp.* strain UFO1. We also thank Melanie Heinemann for technical assistance. We acknowledge the support by the Federal State of Lower Saxony, Niedersächsisches Vorab CDiff and CDInfect projects (VWZN2889/3215/3266). The funders had no role in study design, data collection and analysis, decision to publish, or preparation of the manuscript.

## Supplementary files

*Figure S1 MSA Viewer output of blastn alignment of CD630-hpdBCA excluding C. difficile.*

*Figure S2 MSA Viewer alignment of the CDIF_cresol_R3 primer region against exemplary non-C. difficile strains.*

*Figure S3 Visualization of nucleotide differences in representative detection sequences of specific C. difficile MLST/clade genomes.*

*Figure S4 Top 30 genera in all environmental samples based on 16S rRNA gene sequencing analysis.*

*Figure S5a Bacterial community composition of all environmental samples.*

*Figure S5b Taxonomy legend to Fig. S5a.*

*Figure S6 Alpha diversity as ASV richness of all environmental sample. Table S1: Details of environmental samples*

*Table S2. Information of complete, unique C. difficile genomes available on NCBI on 30 August 2022.*

*Table S3: Output of whole-genome ANIm (wgANIm, Fig. 1); used for calculations of clade-wise wgANIm comparisons (Fig. 2).*

*Table S4. Output of detection sequence ANIm (detANIm, Fig. 3); used for calculations of clade-wise detANIm comparisons (Fig. 2).*

*Data file S1. Multifasta of the 139 detection sequences used in detANIm analysis, with corresponding GenBank accession and MLST information*

*Data file S2. Raw 16S rRNA gene sequencing ASV table Data file S3. Raw 16S rRNA gene sequencing ASV table of amplicons ran with 35 PCR cycles*

